# Adaptation to heavy-metal contaminated environments proceeds via selection on pre-existing genetic variation

**DOI:** 10.1101/029900

**Authors:** Kevin M. Wright, Allison Gaudinier, Uffe Hellsten, Annie L. Jeong, Avinash Sreedasyam, Srinidhi Holalu, Miguel Flores Vegara, Arianti Rojas Carvajal, Chenling Xu, Jarrod A. Chapman, Robert Franks, Jane Grimwood, Kerrie Barry, Jerry Jenkins, John Lovell, Graham Coop, Jeremy Schmutz, John K. Kelly, Daniel S. Rokhsar, Benjamin K. Blackman, John H. Willis

**Affiliations:** Department of Biology, Duke University, Durham, NC, USA; Actio Biosciences, San Diego, CA, USA; Department of Plant and Microbial Biology, University of California, Berkeley; Department of Molecular and Cell Biology, University of California, Berkeley, CA, USA; Genome Sequencing Center, HudsonAlpha Institute for Biotechnology, Huntsville, AL, USA; US Department of Energy Joint Genome Institute, Berkeley, CA, USA; Department of Plant and Microbial Biology, North Carolina State University; Department of Evolution and Ecology, University of California, Davis; Center for Population Biology, University of California, Davis; Department of Ecology and Evolutionary Biology, University of Kansas, Lawrence; Chan Zuckerberg Biohub, San Francisco, CA, USA; Molecular Genetics Unit, Okinawa Institute of Science and Technology, Onna, Japan; Department of Integrative Biology, University of California, Berkeley

## Abstract

Anthropogenic environmental changes create evolutionary pressures on populations to adapt to novel stresses. It is as yet unclear, when populations respond to these selective pressures, the extent to which this results in convergent genetic evolution and whether convergence is due to independent mutations or shared ancestral variation. We address these questions using a classic example of adaptation by natural selection by investigating the rapid colonization of the plant species *Mimulus guttatus* to copper contaminated soils. We use field-based reciprocal transplant experiments to demonstrate that mine alleles at a major copper tolerance locus, Tol1, are strongly selected in the mine environment. We assemble the genome of a mine adapted genotype and identify regions of this genome in tight genetic linkage to Tol1. We discover a set of a multicopper oxidase genes that are genetically linked to Tol1 and exhibit large differences in expression between tolerant and non-tolerant genotypes. We overexpressed this gene in *M. guttatus* and *A. thaliana* and found the introduced gene contributes to enhanced copper tolerance. We identify convergent adaptation loci that are additional to Tol1 by measuring genome-wide differences in allele frequency between pairs of mine and off-mine populations and narrow these regions to specific candidate genes using differences in protein sequence and gene expression. Furthermore, patterns of genetic variation at the two most differentiated candidate loci are consistent with selection acting upon alleles that predates the existence of the copper mine habitat. These results suggest that adaptation to the mine habitat occurred via selection on ancestral variation, rather than independent *de novo* mutations or migration between populations.

## Introduction

Human actions have frequently caused abrupt and severe environmental changes by destroying habitats, changing hydrological regimes, altering community compositions, introducing toxic pollutants and chemicals, and dispersing pathogens, predators, and competitors. Some species have rapidly adapted to survive, or even thrive, when consistently confronted with such novel selective regimes over just tens to hundreds of generations(*1–5*). These classic examples of evolution in action reveal how the genetic diversity adaptive to human-altered environments arises, and inform what conditions enable or constrain the likelihood and speed of evolutionary rescue (*6–8*). Importantly, in many of these cases, adaptive alleles can be aged relative to a known onset of environmental change. Consequently, these examples provide valuable systems for assessing how important different types of mutations and how consequential different types of hard or soft selective sweeps from *de novo* mutations, standing variation, or introgressed variants are in promoting rapid, polygenic adaptation under different demographic scenarios (*9–12*).

Since pioneering studies in the 1950s, plant species inhabiting soils polluted by extreme concentrations of heavy metals, such as mine tailings, have been excellent model systems for investigating the physiological and genetic basis of rapid adaptation to human-altered environments (*13*, *14*). Although quantitative genetic studies have indicated that adaptation to mine habitats is polygenic and some physiological mechanisms that contribute to heavy metal tolerance of mine populations have been described (*15–17*) the specific molecular basis of recent adaptation to any mine environment is not known. Consequently, although some genetic changes have been identified that allowed plants to adapt to soils naturally contaminated by heavy metals over much longer timescales have been identified (*18*), it remains unknown whether similar functional and evolutionary mechanisms can and do rapidly adapt populations to newly exposed toxic soils.

Here, we investigate how populations of the common monkeyflower (*Mimulus guttatus syn. Erythranthe guttata*) have rapidly adapted to thrive on the novel environmental stress imposed by highly toxic copper mine tailings. Copper is an essential micronutrient; however, excess copper inhibits plant growth (*19*). Copper mining experienced a boom in the California Sierra Nevada foothills in the 1860s due to rich veins of copper ore in and around the present-day and aptly named town of Copperopolis in Calaveras County (*20*). Although heavy mining activity only lasted for about five years, the landscape surrounding the mines remains dramatically altered. Nonetheless, large *M. guttatus* populations thrive in vernal streams that run across the otherwise barren surfaces of excavated mine tailings at five sites in Calaveras County (*21*). Previous work on this system notably demonstrated that local adaptation can drive the evolution of reproductive isolation as a byproduct through genetic hitchhiking (*22*, *23*). Initial genetic crossing studies that imposed stress in hydroponic assays determined that copper tolerance is conferred by dominant allele at a single major-effect locus, *Tolerant1* (*Tol1*) (*15*, *24*). Heterogeneity in the degree of tolerance within mine populations also suggested that an undetermined number of additional loci may also contribute to tolerance (*21*). However, the relative contribution of the *Tol1* locus to fitness in mine tailing and off-mine environments has not been assessed, and the molecular identities and evolutionary origins of adaptive allelic variants at *Tol1* and any additional loci contributing copper tolerance remain unknown. Here, we show that rapid adaptation to the mine environments evolved in one part through recent gene duplication at *Tol1* and in another part via selection on ancestral alleles that pre-date the mine environment at multiple other loci.

## Results

### The *Tol1* locus impacts fitness in the mine habitat

To determine whether tolerant alleles at *Tol1* confer local adaptation to copper-contaminated mine tailings, we conducted a reciprocal transplant field experiment using near-isogenic lines (NILs). We employed marker-assisted-backcrossing to breed NILs that were homozygous at *Tol1* for either the mine allele (CCC; Copperopolis, CA) or an off-mine allele (MED; Moccasin, CA). Lines were either backcrossed to CCC inbred lines for four generations (lines: *BC_4_^CCC^_Tol1^CCC^* and *BC_4_^CCC^_Tol1^MED^*) or MED for a single generation (lines: *BC_1_^MED^_Tol1^CCC^* and *BC_1_^MED^_Tol1^MED^*) (Supplemental Figure 1). To remove the effects of inbreeding depression, we backcrossed lines to two different CCC or MED inbred lines per genomic background and then intercrossed those lines to produce outbred progeny (Supplemental Figure 1). In addition to the four focal NIL genotypes, we included outbred CCC and MED parental lines and CCC x MED F_1_s in the experiment. In total, we planted 1868 seedlings into two experimental plots at the defunct Napoleon copper mine (Telegraph City, CA) and at two off-mine, uncontaminated plots located within 2km of the mine site (Figure 1A, B) (Supplemental Table 1).

**Figure 1:**
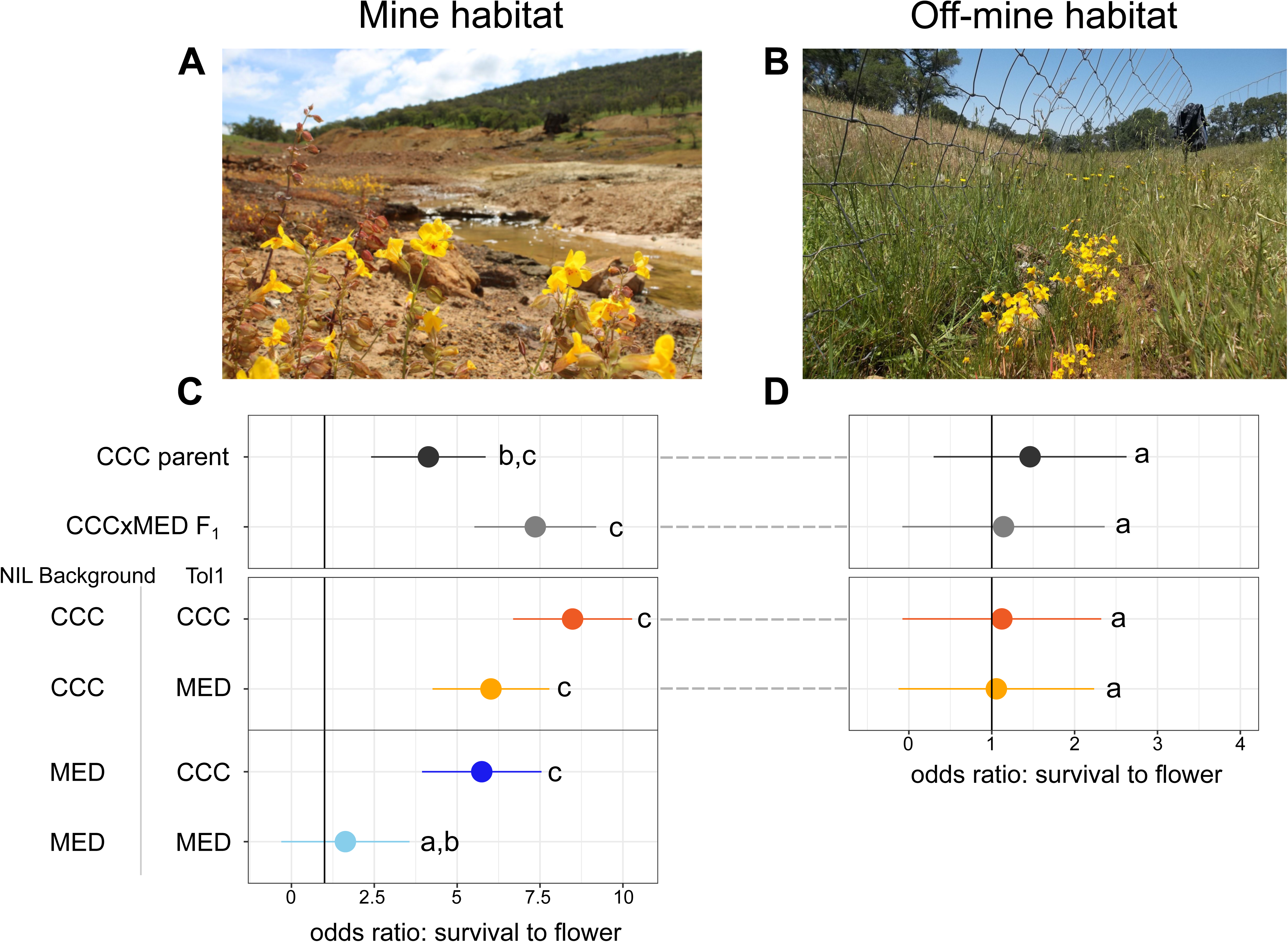
Tol1 locus and background genotype affect survival on copper contaminated mine soil. **A.** Image of *M. guttatus* growing in copper contaminated soil adjacent to mine experimental plots at the Napoleon Mine. **B.** Image of *M. guttatus* growing in uncontaminated soil at an off-mine experimental plot. **C.** Survival analysis for mine habitat. **D.** Survival analysis for off-mine habitat. **C. and D.** Odds of survival to flower are plotted as a ratio of the focal line listed along y axis versus the off-mine, non-tolerant parental line: MED. NILs were bred by backcrossing to either the MED or CCC (mine, tolerant parent) genomic backgrounds. NILs for each genomic background were homozygous for CCC or MED alleles at Tol1. Odds ratios with notation *a* were not significantly different from MED parent, fdr adjusted *P* > 0.05. Notations *b* and *c* denote groups that were significantly different from MED parent and each other at fdr adjusted *P* < 0.05.

We found the CCC parental line was significantly more likely to survive to flower than the MED parental line in the mine habitat (Figure 1C; odds ratio CCC relative to MED = 4.13, fdr adjusted *P* = 0.009). Consistent with the dominant effect of *Tol1* on copper tolerance in lab conditions (*21*), the CCC x MED F_1_ plants were also significantly more likely to survive to flower than the MED plants in the mine habitat (Figure 1C). Introduction of the CCC allele at *Tol1* into the non-tolerant MED genomic background (line: *BC_1_^MED^_Tol1^CCC^*) significantly increased the probability of survival to flower relative to the MED parent (Figure 1A; odds ratio = 5.74, *P*_adj_ =0.003). This result contrasts with the lack of increased survivorship for the MED allele at *Tol1* in the MED genomic background (line: *BC_1_^MED^_Tol1^MED^*) relative to the MED parent (Figure 1C; odds ratio = 1.62, *P*_adj_=0.46). NILs with the CCC background, regardless of *Tol1* allele, *BC_4_^CCC^_Tol1^CCC^* and *BC_4_^CCC^_Tol1^MED^*, possessed significantly higher odds of survival to flowering compared to the MED parent (Figure 1C). The *BC_4_^CCC^_Tol1^CCC^* lines also had an increased probability of survival to flower compared to *BC_4_^CCC^_Tol1^MED^*, but this increase was not statistically significant (Figure 1C; odds ratio = 1.40, *P*_adj_ =0.24). Notably, allelic variation at loci besides *Tol1* affect fitness in the mine habitat, as the CCC genomic background significantly increased survivorship to flower compared to the MED genomic background for NILs with MED allele at Tol1 (Figure 1C; *BC_4_^CCC^_Tol1^MED^* and *BC_1_^MED^_Tol1^MED^* - odds ratio = 3.67, *P*_adj_ =0.007). Together, our results demonstrate that the mine allele at *Tol1* is sufficient to increase survivorship on copper mine tailings, but an unknown number of additional genetic factors are also involved in adaptation to this habitat.

In the off-mine habitat, the parental and CCC x MED F_1_ genotypes did not differ in the likelihood of survival to flower in the off-mine habitat (Figure 1D). Additionally, there was no significant effect of the *Tol1* locus in the CCC background in the off-mine habitat (Figure 1D). Thus, we did not observe a cost to the *Tol1* mine allele in the off-mine environment.

### Triplicated multi-copper oxidase gene maps to *Tol1*

Previously, we introgressed the dominant copper tolerant allele from CCC, *Tol1^CCC^*, into the non-tolerant background of the STB (Stinson Beach, CA) population by seven generations of backcrossing and scoring of copper tolerance (*23*). Among the phenotyped progeny, one heterozygous BC_7_*^STB^ Tol1^CCC^/tol1^STB^* plant was selected and crossed to a homozygous BC_7_*^STB^ tol1^STB/STB^* sibling to generate a mapping population of 4,340 progeny. Using molecular markers and the initial *Mimulus guttatus* v1.0 draft genome assembly of the Oregon inbred line IM62 (*25*) we identified flanking markers to map the *Tol1* locus to a non-recombining, poorly assembled pericentromeric region in the center of chromosome 9. With our most tightly linked marker at 0.32 cM, we identified the recombinant NIL, 25_E11, with the smallest introgression interval but remained heterozygous at *Tol1*.

Next, we identified the location of the *Tol1* flanking markers in a recently generated chromosome-scale assembly of *Mimulus guttatus* var. IM62 (Figure 2A) (*26*). These markers flank what appears to be a contiguously assembled genomic region spanning 1.6 Mbp and containing 48 annotated genes. To identify which, if any, of these genes may be *Tol1* we used the 25_E11 NIL to generate a new fine-mapping population. We selfed the original *Tol1^CCC^/tol1^STB^*25_E11 line and used the tightly linked *Tol1* marker to identify sets of progeny that were homozygous for the copper tolerant allele, *Tol1^CCC/CCC^*, or homozygous for the non-tolerant allele, *tol1^STB/STB^*. To identify candidate genes, we leveraged the alternative homozygous progeny in three complementary experiments.

**Figure 2:**
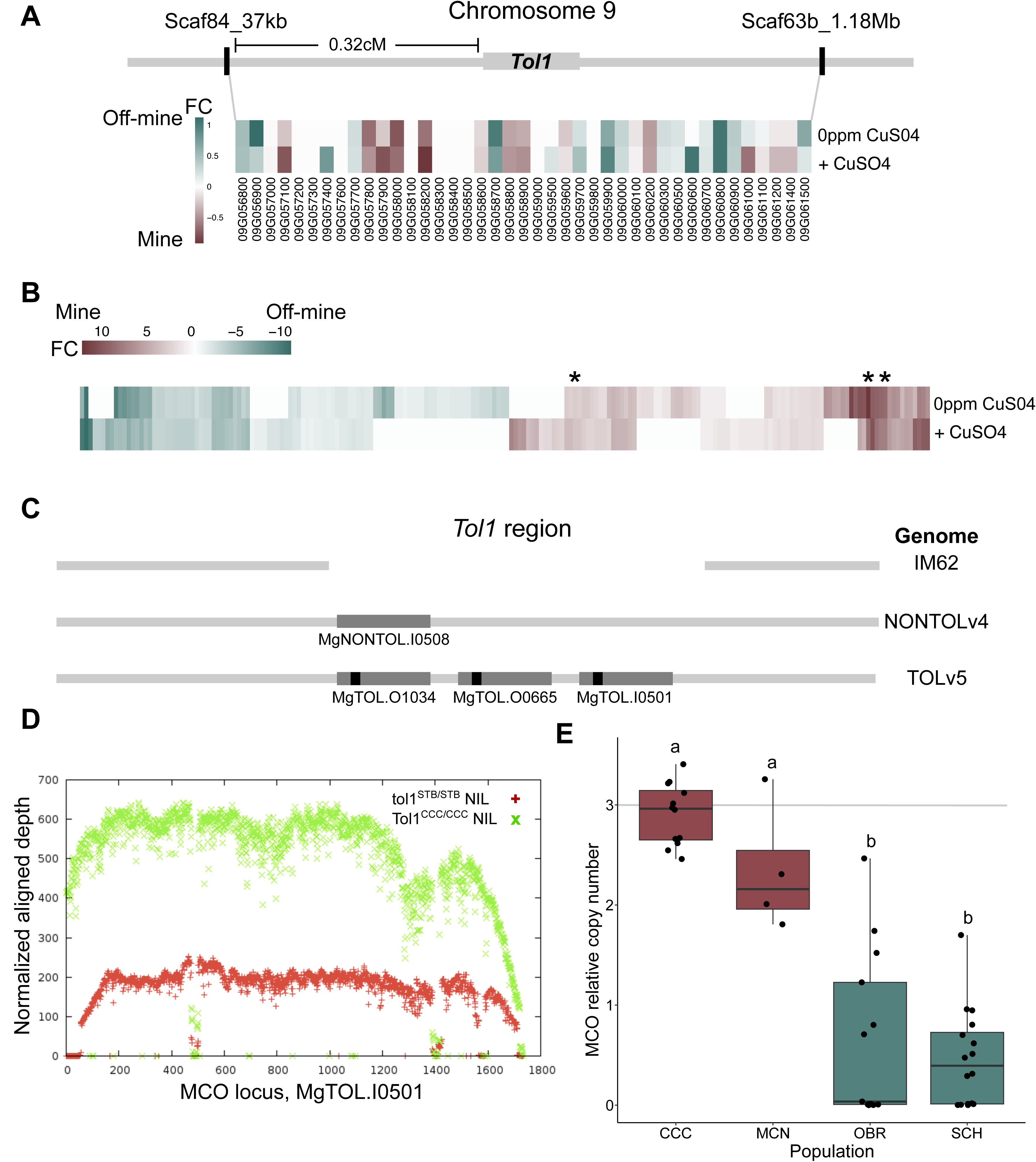
MCO is a candidate gene in Tol1 and has undergone copy number expansion in mine populations. **A.** Schematic of *Tol1* region in Chromosome 9 and gene expression heatmap. Gene expression was measured in *Tol1^CCC/CCC^* and *tol1^STB/STB^* genotypes grown in control and +CuSO_4_ and aligned to IM62 v3.1 genome. Heatmap includes all annotated genes that were located between *Tol1* markers in the IM62 v3.1 genome. No genes were significantly differentially expressed between genotypes in either growth condition (fdr adjusted *P* > 0.05). **B.** Heatmap of all differentially expressed transcripts between *Tol1^CCC/CCC^* and *tol1^STB/STB^*genotypes when mapped to the *Tol1^CCC^/tol1^STB^ de novo* transcriptome. The three MCO transcripts are indicated with *. As with A, genotypes grown in control and +CuSO_4_. All genes displayed exhibited significant differences in fold change and expression: log2 fold change > 1.0 fdr adjusted *P* < 0.05. **C.** Schematic displaying copy number variation for MCO genes in three fully assembled *M. guttatus* genomes: 1) IM62: a highly diverged, non tolerant line from Oregon, 2) NONTOLv4: the non-tolerant NIL *tol1^STB/STB^*, and 3) TOLv5: the tolerant NIL *Tol1^CCC/CCC^*. **D.** Depth of coverage at MCO locus, MgTOL.I0501, for sequenced NILs Tol1 CCC/CCC (green x) and tol1STB/STB (red +). Reads aligned to an edited version of *M. guttatus* TOLv5 genome that possess only a single MCO gene copy. **E.** qRT-PCR measures MCO copy number in wild-collected lines from mine (CCC and MCN) and off-mine (OBR and SCH) populations. Copy number is displayed relative to the tolerant NIL *Tol1^CCC/CCC^*with 3 copies.

First, reasoning that other known examples of heavy metal tolerance in plants often involve increased expression of tolerance-enhancing genes (*16*, *27*), we performed an RNAseq experiment to identify any genes differentially expressed between the alternative homozygous 25_E11 NIL progeny genotypes. We grew replicates of both the *Tol1^CCC/CCC^* and *tol1^STB/STB^* genotypes in either control or CuSO_4_ enriched hydroponic conditions and sequenced RNA extracted from roots (Supplemental Table 3). Of the 41 genes in the Tol1 interval with measurable gene expression, none were significantly differentially expressed with an fdr adjusted *P* < 0.05 and a log_2_ fold change > 1.0 (Figure 2A, Supplemental Table 4). To explore the possibility that expressed transcripts in the NILs may not exist in the IM62 reference genome, we constructed a *de novo* transcriptome from this RNAseq dataset and then recalculated differences in gene expression. This analysis revealed 200 transcripts that were differentially expressed (fdr adjusted p--value < 0.05 and log_2_ fold change > 1.0) (Figure 2C, Supplemental Table 5). In this list, the presence of three distinct transcripts annotated as multicopper oxidase (MCO) genes caught our attention because they were among the highest differentially expressed genes between the genotypes (rank 1: 12.6X fold change, padj<2.5e-17; rank 9: 9.29X fold change, padj<1.1e-37; rank 52: 3.17X fold change, padj<2.1e-21), and none of these transcripts were present in the IM62 reference genome. Enzymes in the MCO superfamily are capable of oxidizing a wide range of substrates while reducing oxygen to water (*28*). Some MCOs have been shown to maintain copper homeostasis by oxidizing Cu(I) to less toxic Cu(II) in bacteria and fungi tolerant to toxic copper levels (*29–31*).

Second, we sequenced the sets of alternative homozygous 25_E11 progeny:*Tol1^CCC/CCC^* and *tol1^STB/STB^* and constructed two new genome assemblies: *Mimulus guttatus* TOL v3.1 and *Mimulus guttatus* NONTOL v4.0. These genomes allowed us to ask whether any differentially expressed genes (including the MCO-like genes) exhibit presence-absence variation within the *Tol1* region between the Oregon inbred line used for the *M. guttatus* var. IM62 v3.1 genome and the California CCC and STB populations. In both the NONTOL v4.0 and TOL v3.1 genomes we found a single copy of the differentially expressed MCO-like genes (MgNONTOL.I0508, MgTOL.I0501) in their chromosome 9 assemblies. In addition, two additional copies of the MCO-like gene (MgTOL.O0665 and MgTOL.O1034) were present on two small scaffolds that were not incorporated into any chromosomes of the TOL v3.1 assembly.

Third, using the many plants of each of the alternative 25_E11 NIL progeny genotypes *Tol1^CCC/CCC^*and *tol1^STB/STB^*, we performed bulked segregant analysis to determine whether those additional copies of the MCO-like gene in the *Tol1^CCC/CCC^* genotype also map to the *Tol1* locus (Supplemental Figure 2). To assess whether all three MCO-like genes were linked to Tol1, we compared the depth of coverage at this locus between the *Tol1^CCC/CCC^*and *tol1^STB/STB^* bulk pools. We found a nearly 3x difference in depth of coverage between these pools (Figure 2D). This result suggests all three MCO-like genes are located within the *Tol1* introgression interval.

Taken together, through comparison of *Tol1^CCC/CCC^* and *tol1^STB/STB^* NIL progeny, we find that the tolerant *Tol1^CCC^* allele contains three differentially expressed copies of an MCO-like gene, whereas the non-tolerant *tol1^STB^* allele contains only one copy, and surprisingly this MCO-like gene is completely absent from the entire Mimulus guttatus var. IM62 reference genome.

### Elevated MCO copy number is pervasive in mine and nearly absent from off-mine plants

To determine whether the MCO triplication in *Tol1^CCC/CCC^* plants was abundant in mine populations and not just a difference between the specific NILs used for the previous experiments, we measured MCO copy number variation in wild collected mine and off-mine plants using quantitative-PCR. We found the median MCO copy number to be 3.0 and 2.2 at two mine populations (Figure 2E, Supplemental Table 6). In contrast, MCO copy number was significantly lower in the two offline populations, with median copy number values equal to 0.5 and 0.1 (Figure 2E). To further examine natural variation in MCO copy number, we used genome re-sequencing data from a broad collection of plants sampled from mine tailings or local off-mine sites to measure the number of reads that aligned to any of the three MCO gene copies in the TOL v3.1 genome assembly in comparison to non-MCO genes located within 1Mb. As expected, we found excess read coverage at MCO in mine plants compared to off-mine plants, but we did not observe a similar pattern at adjacent genes (Supplemental Figure 3). Thus, MCO copy number is elevated in all mine plants and low or absent in nearly all off-mine plants, consistent with past observations that tolerance is nearly fixed in mine populations and segregates at very low frequencies in adjacent offline populations (*21*).

### *MgMCO* overexpression improves copper tolerance of non-tolerant genotypes

To test the hypothesis that the higher MCO expression levels, potentially resulting from copy number expansion, confers increased tolerance in copper mine populations, we asked whether overexpression of an MCO copy from a Cu tolerant population (CCC52 MgTOL.O0665) is sufficient to increase copper tolerance in non-tolerant genetic backgrounds. We transformed a 35S:MCO construct into wild-type *M. guttatus* (MAC) and *Arabidopsis thaliana* (Col-0). We assayed mutant seedlings for root growth phenotypes as a proxy for tolerance on plates containing excess CuSO_4_. In the *M. guttatus* mutant lines we found that the primary roots of the 35S:MCO lines grew significantly longer by 0.26-0.93 cm following transfer to plates supplemented with 20ppm (80μM) CuSO_4_ compared to WT MAC seedlings (one-way ANOVA *P* < 0.05, Supplemental Table 7a). Similarly, the *A. thaliana* Col-0 background, the primary roots of 35S:MCO lines grew significantly longer by 0.93-2.22 cm compared to WT plants when grown on media supplemented with 15ppm (60μM) CuSO_4_ (one-way ANOVA *P* < 0.05, Supplemental Table 7b). Thus, higher expression levels of MCO, associated with the copy number expansion of this gene in the mine populations, likely contribute to the enhanced copper tolerance conferred by the *Tol1* locus.

### Candidate mine habitat adaptation loci identified from selective sweeps and differential expression

Although we increased copper tolerance of off-mine genetic backgrounds through introgression of the tolerant allele at major locus *Tol1* and by overexpressing a mine copy of MCO, neither intervention yielded plants as tolerant as mine *M. guttatus* populations. Therefore, adaptation to the mine tailing habitat must involve additional loci. To identify these loci, we performed whole genome sequencing of pools of population samples from two pairs of mine and off-mine populations (Pair 1: CCC and OBR; Pair 2: MCN and SCH). For each pair, the genome-wide average value of genetic differentiation measured by F_ST_ ranged from 0.050 to 0.079, suggesting high levels of gene flow or a short time since divergence between these geographically neighboring populations (Supplemental Table 2). To identify regions of convergent differentiation between mine and off-mine adapted populations, we conducted a genome-wide scan for parallel selective sweeps using the likelihood ratio test statistic: LRT-1 (*32*). We detected 16 genomic regions each spanning between two and twenty-one genes with signatures of parallel selection (Figure 4A, Supplemental Table 8).

**Figure 3:**
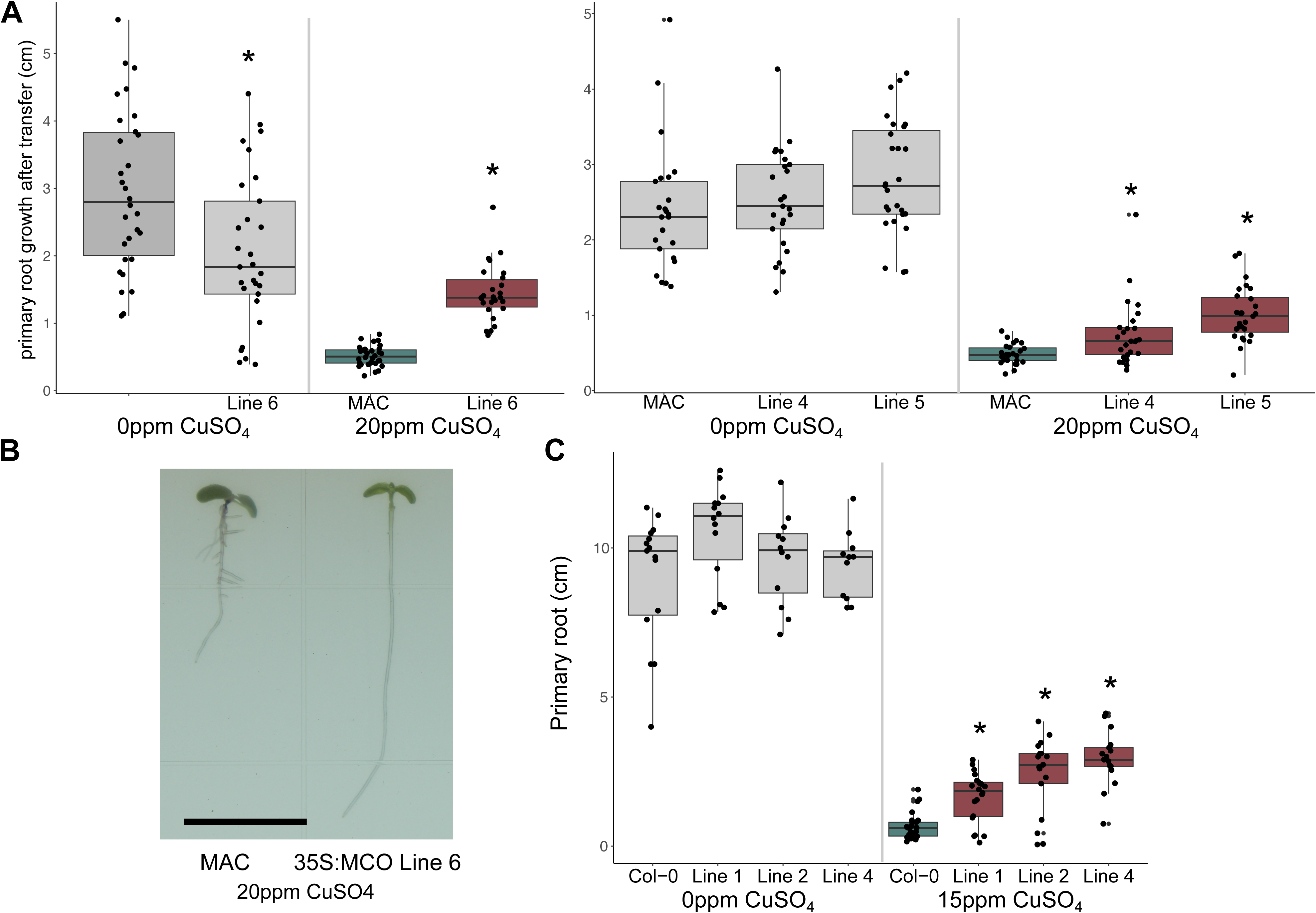
35S:MCO transgenics tolerate excessive Cu stress. **A.** M. guttatus 35S:MCO lines grown in control and 20ppm CuSO_4_ compared to parental control (MAC). All lines shown are significant. Color reflects tolerance (teal = non-tolerant, maroon = increased tolerance). * indicates *P* < 0.05 using a one-way ANOVA compared. **B.** Image of M. guttatus MAC and 35S:MCO Line 6 grown in 20ppm CuSO_4_. Black bar = 1cm **C.** A. thaliana 35S:MCO lines grown in control and 15ppm CuSO_4_ compared to parental control (Col-0). All lines shown are significant. (teal = non-tolerant, maroon = increased tolerance). * indicates *P* < 0.05 using a one-way ANOVA.

**Figure 4:**
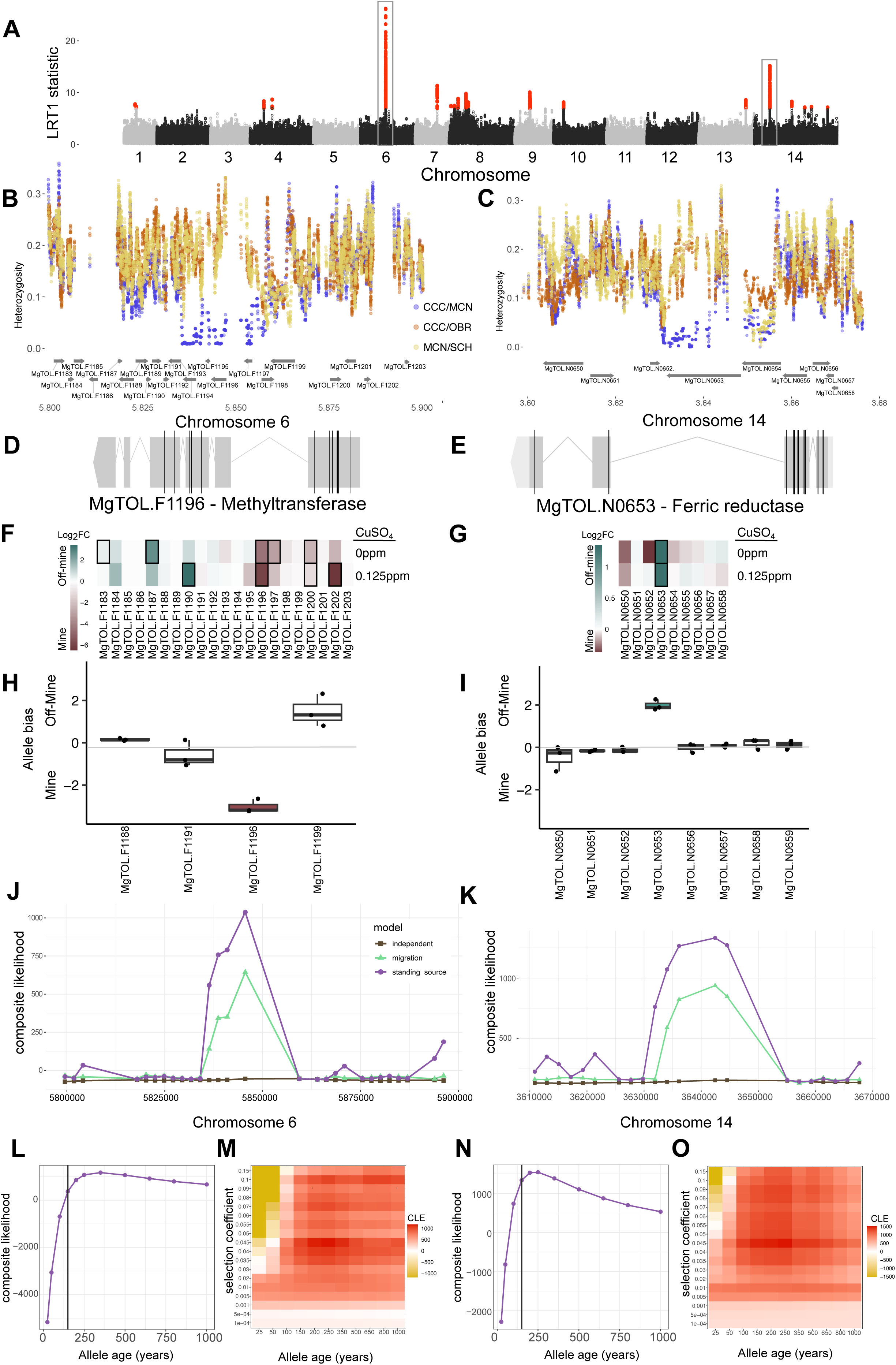
Additional loci show signatures of selection on copper mines from standing variation. **A.** Genome-wide plot of LRT-1 values depict regions of parallel adaptation in mine populations. Each point is the average of 30 adjacent SNPs. Red colored points highlight SNP bins in the 0.01% tail of LRT-1 values. **B-C**. Heterozygosity plots for candidate adaptation loci on chromosome 6 and 14. Heterozygosity between populations was measured for: CCC(mine) and MCN(mine) in blue, CCC and OBR(offmine) in orange, MCN and SCH(offmine) in gold. **D-E**. Gene plot displaying SNPs for MgTOL.F1196 (methyltransferase) and MgTOL.N0563 (ferric reductase) segregating between mine and off mine genomes. Black bars indicate nonsynonymous SNPs. **F-G**. Heatmap showing differential expression for candidate adaptation genes in chromosome 6 and chromosome 14, respectively. Genes that are significantly differentially expressed (fdr adjusted *P* <0.05) and have a log2 fold change greater than 1.0 are surrounded by a black box. Higher expression in off-mine parent is teal, higher expression in mine parent is maroon. **H - I**. Allele specific expression analysis for candidate adaptation genes in chromosome 6 and chromosome 14, respectively. Points are biological replicates, with the 0.25 to 0.75 quartile range shown in a box plot. Teal colored boxes are significantly biased toward off-mine parent, maroon colored boxes are significantly biased toward mine parent (t-test, *P* < 0.05). Open box plots showed no evidence of bias. **J - K.** DMC model compares three sources of adaptive allelic variation at the chromosome 6 and 14 loci: independent *de novo* mutation (grey), recent migration (green), and ancient standing variation (purple). The x-axis depicts the genomic intervals for which the models were evaluated. For each locus, we split the region into 100 bins equally spaced on physical distance. Bins were filtered if they contained less than 20% of the maximum number of SNPs per bin per locus. **L, N.** Composite likelihood estimate of age for the selected allele at chromosomes 6 and 14 from standing variation DMC model. Allele age units are years, which are equal to generations for this annual plant population. **M, O.** Pairwise composite likelihood estimate of age and selection intensity for the selected allele at chromosomes 6 and 14 from standing variation DMC model. Heatmap color represents composite likelihood estimate: CLE.

Because each parallel selective sweep region contained multiple genes, we evaluated several criteria to identify candidates with the strongest support. First, we determined which gene(s) have the lowest heterozygosity measured between the two mine populations (CCC and MCN) relative to heterozygosity measured between the paired mine/off-mine populations. This pattern is consistent with a single shared haplotype at high frequency in CCC and MCN, that is distinct from the haplotypes segregating in either off-mine population. Second, we determined whether any SNPs with a significant LRT-1 statistic (*P* < 1e-7) were nonsynonymous as potential indicators of candidate genes that may have contributed to mine adaptation due to changes in protein function. Third, we measured gene expression for individuals from a mine parental genotype, from an off-mine parental genotype, and their F_1_ hybrid line when grown in control or copper enriched hydroponic solutions. We tested for differential expression between the parental genotypes to reveal candidate genes that may have contributed to mine adaptation due to changes in gene expression. Moreover, we assessed patterns of allele-specific expression in the F_1_ hybrid line to determine whether the differences in gene expression between the parental genotypes are due to *cis*-acting regulatory variants that are likely to be within the selective sweep region.

Of the selective sweep regions, six have a genomic interval of at least 1kb with a shared haplotype (pi_CCC_MCN_<0.1) in the mine populations that is highly differentiated from the off-mine populations (pi_CCC_MCN_-pi_CCC_OBR_ >0.10 and pi_CCC_MCN_-pi_MCN_SCH_ >0.10). For the genes that displayed low heterozygosity within the 16 sweep regions in the mine populations, we analyzed 72 genes for SNPs: 66 genes had no significant SNPs with elevated LRT-1 values that cause nonsynonymous substitutions and the remaining six genes had one to 15 nonsynonymous substitutions (*P* < 1e-7, Supplemental Table 9). Of the 179 total annotated genes within the parallel sweep regions, 56 (31%) were significantly differentially expressed (log_2_FC > 1, *P_adj_* < 0.05) between the tolerant and non-tolerant parental genotypes grown in either control or copper-enriched (+Cu) conditions (Supplemental Table 10). Interestingly, no genes exhibited a significant genotype-by-environment interaction effect. Of the 66 expressed genes in the sweep regions that could be differentiated by parental SNPs, we found allelic bias for 12 genes (18%) in control conditions and/or +Cu conditions in the F_1_ hybrids. We found one or two genes in 11 of the sweep regions that had allelic bias in their expression (Supplemental Table 11).

Applying these criteria highlighted several strong candidate genes for copper adaptation. For instance, the largest LRT-1 peak in the genome resides on chromosome 6 and encompasses 21 genes (Figure 4A, B, Supplemental Table 8). Examining shared heterozygosity between mine populations and reduced heterozygosity relative to offline populations reduced the region of interest to five candidate genes: MgTOL.F1194 - MgTOL.F1198 (Fig. 4B). Of these candidates, two–MgTOL.F1194, a FAR1-related sequence protein, and MgTOL.F1196, a methyltransferase– had five and twelve variants with elevated LRT-1 values that cause non-synonymous substitutions, respectively (Figure 4D, Supplemental Table 9). MgTOL.F1196 was expressed at significantly higher levels in mine plants relative of off-mine plants in both the control (log_2_FC of 4.5) and +Cu (log_2_FC of 6.1) conditions; one other annotation, MgTOL.F1197, was significantly differentially expressed in the control conditions (log_2_FC of 3) (Figure 4F). Of these two genes, only MgTOL.F1196 showed evidence of allele-specific expression in the F_1_ hybrids (Figure 4F). These results suggest that the methyltransferase MgTOL.F1196 is the best supported candidate gene for mine adaptation in this region. Notably, homologs of this gene have not been previously associated with copper homeostasis in plants.

The second largest LRT-1 peak in the genome resides on chromosome 14 and encompasses nine genes (Figure 4A, C, Supplemental Table 8). We found the region where heterozygosity plummets when comparing the two mine populations relative to other comparisons includes two genes, MgTOL.N0653 and MgTOL.N0654 (Fig. 4C). MgTOL.N0653, a ferric reductase, bears 15 SNPs differentiating mine and off-mine populations that cause non-synonymous substitutions while MgTOL.N0654 bears none (Figure 4E, Supplemental Table 9). Of the nine genes in this LRT-1 peak, only MgTOL.N0653 exhibited significant differential expression (log_2_FC of 1.4 and 1.3 in control and +Cu conditions) Figure 4G, Supplemental Table 10), and this differential expression was attributable to *cis*-acting regulatory differences (Figure 4I, Supplemental Table 11). These results are consistent with MgTOL.N0653 being the strongest candidate in this sweep region.

The functional effect of MCO on copper tolerance as well as the geographical distribution of this gene: high copy number in all mine adapted populations that we have investigated and absent from all other *M. guttatus* populations except a few located near Copperopolis, CA where it is present in only a single copy, suggests this locus may exhibit strong patterns of a parallel selective sweep. However, when we applied these same criteria to the MCO triplicated gene locus, we did not find evidence of parallel selective sweep. This locus is not among the top 16 loci highlighted in genome-wide LRT-1 analysis (Supplemental Figure 4A), although a few individual SNPs in this region have LRT-1 p-values < 1e-7 (Supplemental Figure 4B). Heterozygosity between CCC-MCN mine populations is only slightly reduced in comparison to CCC-OBR or MCN-SCH at a single one MCO copy MgTOL.O0665, and showed weaker differences at the remaining two loci (Supplemental Figure 4C, D). Though there was no conclusive evidence of a parallel selective sweep MCO, we suspect this test was not as well suited for this MCO locus (in comparison to other candidate loci) because we have observed variation in MCO copy number segregating in mine populations (Figures 2E, S Figure 3) and this may artificially inflate heterozygosity values in the mine populations and decrease the signal of a recent selective sweep.

Although homologs of the ferric reductase and some other candidates in the parallel selective sweep regions have been previously implicated in copper homeostasis (*33*) (Supplemental Table 12), no function has been attributed to the homolog of the methyltransferase and other top candidate genes (Supplemental Table 12). Thus, by studying naturally occurring, adaptive allelic variation, our work has highlighted new targets for future mechanistic work aimed at understanding how plants maintain balanced levels of this essential micronutrient.

### Mine adaptation loci evolved via selection on standing variation

To understand the evolutionary process that led to rapid adaptation in the mine environment, we investigated the source of allelic variation for the candidate parallel selection loci identified by the LRT-1 statistic. Three evolutionary scenarios distinguish the origins of the selected alleles: (1) independent mutational origins, (2) single allele shared due to recent migration between populations, or (3) single allele shared due to ancestral segregating variation. We applied the DMC likelihood model to analyze the distribution of genetic diversity between populations at our two lead candidates to distinguish the likelihood of different sources of adaptive alleles as well as estimate the selection coefficient and the age of these alleles (*34*).

We found that for the two lead candidate adaptation loci, the standing variation model had the greatest likelihood score (Figure 4J, K). The standing variation model is consistent with the observation of a single haplotype that is shared between the two mine populations at the core of the selected locus (Figure 4B, C). In contrast, we do not find support for independent sweeps of distinct mutations in each population, because this would cause high genetic differentiation between the two mine populations at the core of the selected locus. Additionally, the data does not support recent migration between habitats because that would cause the shared haplotype to extend over a much larger genomic region.

Next, we estimated the age of the selected alleles using the rate at which genetic diversity around the mine allele recovers to background levels. The maximum likelihood estimate for the age of the selected alleles at the two lead candidates was 350 years (Figure 4L) and 250 years (Figure 4N) both of which predate the onset of copper mining in the region (150 years). We estimated selection coefficients required for these alleles to reach fixation in the mine populations given their estimated ages to be 0.045 for both loci (Figure 4M, O). The selection coefficient estimates are minimum values and could be much higher if these alleles went to fixation soon after colonization of the mine habitats.

In contrast to the results for chromosome 6 and 14 parallel selection loci, we found that both the recent migration and standing variation models were equally likely to model the distribution of genetic variation between populations at MCO locus (S Figure 4F). We estimated the maximum likelihood estimate for the age of the selected allele to be 50 years (long after copper mining began in the region) and the strength of selection to be 0.10 (S Figure 4G, H). The result that the strength of selection for this locus was double the values estimated for the candidate loci in the most significant LRT-1 peaks was consistent with our field-based measure of strong selection on the mine allele at *Tol1* (Figure 1C). Because the estimated age of the allele post-dates the onset of copper mining and the models assuming recent migration and standing variation cannot be distinguished, we infer that at least some of the adaptive allelic variation at MCO arose by *de novo* mutation to the single copy allele found in nearby off-mine populations and then spread among mine populations.

## Discussion

Together, our results provide strong ecological and genetic evidence that an increase in copy number and expression of a novel MCO gene is one of several changes that have enabled *M. guttatus* populations to rapidly adapt to copper-contaminated mine tailings. Our findings are notably convergent with several other instances of adaptation to heavy metals. At the level of MCO itself, it has been shown that strain-specific gains of a particular multi-copper oxidase gene also confer increased copper tolerance in microbial and fungal systems (*31*, *35*). At the scale of the more general evolutionary mechanism, our findings mirror the well-known triplication and subsequent evolution of a set of HEAVY METAL ATPases that have adapted the zinc hyperaccumulator *Arabidopsis halleri* to heavy-metal rich habitats (*36*, *37*). Notably, our findings indicate such complex, multi-step adaptive genetic changes, which arose within *A. halleri* sometime since its split with *A. lyrata* hundreds of thousands of years ago (*36*), were derived within just a few hundred years in *M. guttatus*.

Two phenomena that may facilitate evolutionary rescue in human-altered environments appear important in our system (*8*). First, subsets of beneficial *de novo* mutations that arise at elevated rates may disproportionately contribute to rapid and parallel adaptation (e.g., (*38*, *39*)), particularly in high fecundity species like *M. guttatus*. Our work adds to the growing number of case studies finding that types of mutation that occur at higher rates than point mutations, like transposable element insertions or gene duplications, often contribute to rapid adaptation to human-altered environments (e.g., (*2*, *4*)). Notably, as appears to be the case for the MCOs, these adaptations can involve multiple such mutations (*40*). In addition to their increased copy number and expression, the *Tol^CCC^* allele’s three copies of MCO, which share >99% protein similarity to each other, are also distinguished from the *Tol^STB^* allele’s single MCO copy by an expansion of simple sequence repeats that add multiple methionines near the N-terminal. Future biochemical studies are needed to determine whether these methionine-rich repeat sequences, which could plausibly evolve rapidly through replication slippage and/or gene conversion, are specifically adaptive and facilitate greater recruitment and detoxification of copper ions, as found for the methionine-rich domain of the relevant MCO protein in copper tolerant strains of *Escherichia coli* (*41*).

Second, our results indicate selection on standing genetic variation at many loci also drove rapid adaptation of *M. guttatus* to copper mine tailings. Notably, although these adaptive alleles pre-existed the California copper mines, they appear to be fewer than 200 generations older than the onset of selection. These findings contrast with many other recently described cases of adaptation from standing genetic variation during the Holocene or just the Anthropocene in which the beneficial alleles have been acquired by adaptive introgression from other taxa (e.g., (*11*, *12*, *31*)) or are much more ancient alleles preserved by balancing selection within species (e.g., (*42–44*)). High effective population sizes, high fecundities, annual life history, and high dispersal rates may all increase the likelihood that soft sweeps of young, rare pre-existing beneficial alleles contribute to evolutionary rescue in human-altered environments (*7*, *8*). Populations of *M. guttatus* can exhibit all of these qualities (*45*, *46*), potentially explaining how this species has rapidly evolved to thrive on such barren and toxic soils.

## Methods

### Section 1. Reciprocal Transplant Experiment

#### Generate Mapping Lines

We conducted a reciprocal transplant experiment using lines with specific genotypes at the TOL1 locus introgressed into mine or non-mine genomic backgrounds. The mine derived parental lines (CCC) were derived from the seeds collected at the central smelter of Copperopolis, CA (Latitude 37.969, Longitude: -120.633) and non-mine parental lines (MED) were from Moccasin, CA (Latitude 37.831, Longitude -120.342). The parental lines were inbred for at least 4 generations prior to breeding the F1 hybrid and near isogenic lines (NILs). We employed marker-assisted-backcrossing to breed NILs that were homozygous for mine alleles (CCC) or non-mine alleles (MED) at the TOL1 locus. Lines were either backcrossed to CCC inbred lines for four generations (lines: *BC4^CCC^_TOL1^CCC^*and *BC4^CCC^_TOL1^MED^* ) or MED for a single generation (lines: *BC1^MED^_TOL1^CCC^* and *BC1^MED^_TOL1^MED^* ) following the crossing design outlined in Supplemental Figure 1). To remove the effects of inbreeding depression, we backcrossed lines to two different inbred parental lines per genomic background and then intercrossed them to produce outbred progeny of each focal genotype (Supplemental Figure 1). Similarly, we used multiple inbred lines to produce outbred progeny for each focal parental and F1 genotype. In total, seven lines were used in the reciprocal transplant experiment: CCC parent, MED parent, CCCxMED F1, and the four focal NIL genotypes in the experiment.

#### Reciprocal Transplant Experiment

The reciprocal transplant experiment was conducted at the Napoleon copper mine (Latitude: 37.927, Longitude: -120.730), located 11 km from Copperopolis, CA. This site possesses a native population of mine adapted *M. guttatus* plants (*47*, *48*). We planted 1868 seedlings into two experimental plots at the Napoleon mine and into two off-mine experimental plots located within 2km of the mine. The off-mine plots were not contaminated with copper and had 100 ppm Cu less than mine sites (Supplemental Material Table 1). All soil testing was performed at the Texas A&M soil testing lab (https://soiltesting.tamu.edu/).

Seeds were planted into potting soil and exposed to ambient weather conditions in January 2012. The soil was kept moist until seeds germinated. All seedlings were transplanted into experimental plots after the production of the first set of true leaves, ∼1-2 weeks post germination. Seedlings were planted into experimental plots with a minimum amount of potting soil. We transplanted seedlings into sites with native *M. guttatus* populations and their exact location was marked with toothpicks. Genotypes were randomly organized in blocks of 12 plants, and 20-40 blocks were planted into each experimental plot. There were two experimental plots for the mine and off-mine habitats. We recorded the exact location of each block within the plot. Due to low germination rates, *BC1^MED^* lines were planted into the mine plots four weeks after the initial planting and none of these lines were planted into the off-mine plots. This February 2012 round of planting was composed of blocks with the following genotypes: parental MED, parental CCC, *BC1^MED^_Tol1^CCC^*and *BC1^MED^_Tol1^MED^*. All seedlings were provided with supplemental water each day for one week following transplant. We censused plants for survival and flowering intermittently from March 1 to April 4 and everyday from April 4 until May 16. No plants in the mine plots survived beyond May 16.

#### Survival Analysis

We calculated the probability of survival to produce at least a single flower using a cox proportional hazard model, implemented in the R package: coxme (*49*). This implementation of the cox proportional hazard model allows for fixed and random effect terms. We assigned fixed effects to be: Tol1 genotype and genomic background. We assigned random effects: to be planting date, experimental plot id and block location within each plot. We ran separate models to estimate survivorship in mine and non-mine habitats. We tested whether the probability of survival to flower was significantly different between all possible pairs of genotypes in the r package: emmeans (*50*). All probability of survival to flower results are presented as an odds ratio with the MED parent as the reference.

### Section 2. Identification of MCO as the Tol1 locus candidate gene

#### Plant Growth and RNA Extraction of Tol1^CCC^ and Tol1^STB^ lines

For this experiment, we used Tol1^CCC^ and Tol1^STB^ lines from the NIL mapping population. We germinated seeds from these lines in potting soil, and transplanted one week old seedlings into hydroponic solution with either 1x Hoagland solution or 1x Hoagland solution + 0.125 ppm CuSO_4_ solution, pH of 7.0. Plants were grown for 20 days, changing the solution every 3 days, in a growth chamber with 8 hours of light and constant temperature of 20°C. We harvested root tissue, extracted RNA using the Zymo Direct-zol RNA miniPrep Kit (Cat. R2070) and purified RNA using the Invitrogen Plant RNA Purification Reagent kit (Cat. 12322-012). We pooled tissue from 4 -14 plants per library. We had three biological replicates for each treatment/genotype for a total of 12 libraries.

#### RNA sequencing and de novo transcriptome assembly

The 12 libraries were sequenced to 130-160X coverage using the Illumina HiSeq 2500 platform at the HudsonAlpha Institute for Biotechnology, Inc., Genomic Services Lab. We processed the RNA-seq reads using custom Python scripts to trim adapter sequences and retain high quality (Q ≥ 25) sequences. We constructed a de novo transcriptome assembly using all high quality reads de novo using Velvet v. 1.2.10 (*51*) and Oases v. 0.2.08 (*52*). We generated multiple K-mer assemblies (47,71,95) and joined contigs using Oases. We merged the transcript assemblies from the Tol1^CCC^ and Tol1^STB^ lines and filtered redundant contigs using cd-hit (*53*). This de-novo transcript contained 221888 non redundant transcripts.

#### Differential Expression Analysis and Annotation

We performed statistical analysis of differentially expressed loci between tolerant and wild type in Copper+ and Hoagland media in R (v3.1.2) using the DESeq2 package (v1.6.3) (*54*). With this experimental design, we conducted four contrasts: 1) TOL1^CCC^ lines control vs. copper hydroponic solution, 2) TOL1^STB^ lines control vs. copper solution, 3) TOL1^CCC^ vs. TOL1^STB^ control solution, and 4) TOL1^CCC^ vs. TOL1^STB^ copper solution. We defined significantly differentially expressed transcripts as those with an Benjamini and Hochberg FDR adjusted p-value less than 0.1 and log2 fold change greater than 1.0. We annotated differentially expressed loci using BLASTX search against NCBI non-redundant protein database with an e-value threshold of 1 x 10-5.

#### Mapping Population Generation

In order to improve the genetic map at Tol1, we created a new mapping population using lines from a Near Isogenic Line (NIL) mapping population that were either homozygous for the Copperopolis allele (Tol1^CCC^) or nontolerant Stinson Beach allele (Tol1^STB^). These lines were derived from selfing a single line, 25E11 with a recombination breakpoint near Tol1 (Wright *et al.* 2013-Figure 2, line 25_E11) and homozygotes were identified by genotyping the offspring from this cross for two markers, *Sc84_37kb* and *Sc84_21kb*, <0.32cM from Tol1. We identified 100 plants from each genotypic class, pooled equal amount of tissue from each plant, and extracted DNA from the two pools. We removed all reads with Phred quality score, Q, < 30. We aligned reads using *bwa* and filtered out all reads with alignment score q < 40. The mean depth of coverage for the Tol1^STB^ pool is 160X and Tol1^CCC^ pool is 170X.

#### BSA Mapping Procedure

To map scaffolds to the TOL1 region we assume that variants in this region should, except in the case of a rare recombination event, be fixed for alternative alleles in the TOL1^CCC^ and TOL1^STB^ pools. We measured recombination rate as the deviation from fixation at informative SNPs along a scaffold. For example, if a site shows 7 *A* and 152 *G* nucleotides in the TOL1^CCC^ pool, but 140 *A* and 5 *G* nucleotides in the TOL1^STB^ pool alignment, we would classify the tolerant variant to be *G* and interpret the *A’s* in this pool to be due to crossovers between this site and the *Sc84_37kb* marker. The maximum likelihood estimate of the recombination distance for a single site would be 7/159 for the TOL1^CCC^ pool, 5/145 for the TOL1^STB^ pool. We define recombination informative SNPs to have 80-600X coverage and the maximum likelihood estimate of recombination is less than 0.10, or 10cM. Using these criteria, we identify 30.2K recombination informative SNPs. We measure the recombination distance between *Sc84_37kb* and a focal genic region, as the median value for the informative SNPs in the region, after discarding outliers beyond 2 standard deviations from the median. Using this dataset, we identified 400 genic regions in all or parts of 8 scaffolds, totaling 4.9 Mb in sequence, with an estimate of recombination less than 0.008, or approximate recombination rate of 0.80 cM.

#### MCO Relative copy number

Quantitative PCR was performed using genomic DNA to determine copy number variation in mine and off-mine populations and Cu tolerant and nontolerant near isogenic lines. CCC (n=7), MCN (n=4), SCH (n=16), OBR (n=13). qPCR was performed on genomic DNA of the inbred parents, CCC9 and MED84. Primers were designed using Primer 3 Plus (*55*) and validated with standard PCR, gel electrophoresis, and shotgun sequencing of the products to ensure the desired segment was amplified. Reactions were prepared using Thermo Fisher Scientific Fast SYBR green master mix (#4385612) and run on a Roche LightCycler 96 per master mix instructions. We used a delta delta CT method to calculate relative quantification of MCO, using RAD51, a known DNA repair gene that has one copy in Mimulus, to normalize values within samples. Results are relativized to the off-mine inbred line. Differences between genotypes was analyzed using a one-way ANOVA using the lm() function in R.

Primers:

RAD51_F GAAGATGGAACCAGCCAGAC RAD51_R CCCGTAAACACCTTTGCAGT MCO_F TGAAGGTCACACCACGAAAA MCO_R CCTCGGATGGAGACAACAAT

### Section 3. Transgenics

#### MCO overexpression construct

MgTOL.O0665 was cloned from the CCC9 background into the pEarlyGate100 plasmid. Sequence confirmed colonies were transformed into *Agrobacterium tumefaciens* strains LBA4404 and EHA105. *Arabidopsis thaliana* Col-0 lines were transformed via the floral dip method with LBA4404. *Mimulus guttatus* MAC lines were transformed via tissue culture with EHA105. Transgenic plants were confirmed through phosphinothricin resistance and PCR validation of transgene.

#### Root phenotyping assays

1. *M. guttatus* and *A. thaliana* 35S:MCO lines were grown with parental controls to eliminate material effects on seedling growth. Modified Hoaglands solution (*56*) was used as the control media: 10 mM KNO_3_, 4 mM MgSO_4_, 2 mM KH_2_PO_4_, 1 mM CaCl_2_, 10 mM KCl, 36 mg/l FeEDTA, 0.146 g/l 2-morpholinoethane sulfonic acid, 1.43 mg/l H_2_BO_3_, 0.905 mg/l MnCl_2_·4H_2_O, 0.055 mg/l ZnCl_2_, 0.025 mg/l CuCl_2_·2H_2_O, 0.0125 Na_2_MoO_4_·2H_2_O, 1% sucrose, 0.75% agar, pH 5.7.
2. *M. guttatus* seeds were sterilized and plated on control media and stratified for one week at 4°C. Plates were transferred to incubators with a 14 hour day/10 hour night cycle in 22°C. Germination of seedlings was recorded and 2 day old seedlings were transferred to new control plates or plates with an additional 20ppm CuSO₄._5_H₂O. Seedlings were imaged on transfer day and one week after transfer.
3. *A. thaliana* seeds were sterilized and plated on control media or media with additional 15ppm CuSO₄._5_H₂O and stratified for 3 days. Plates were transferred to incubators with a 14 hour day/10 hour night cycle in 22°C. Germination of seedlings was recorded and seedlings were imaged at 12 days old.

Primary roots of seedlings were traced using a Wacom tablet in ImageJ. Data were log-transformed and analyzed using a one-way ANOVA using the lmer() function in R (*57*).

### Section 4. Identification and Evolutionary History of Mine Adapted Alleles

#### SNP analysis

72 genes from low heterozygosity regions within the LRT-1 sweeps from CCC, MCN, OBR, and SCH resequenced genomes were analyzed for SNP content. Gene models were based on the *Mimulus guttatus* TOL v5.0 genome. Nonsynonymous SNPs were cataloged and called significant if they had an LRT-1 adjusted p-value < 1E-7.

#### RNA-seq libraries of CCC, MED, and CCCxMED F_1_ in control and +Cu conditions

Seeds were planted on rock wool in cut microcentrifuge tubes in 0.5x Hoagland basal salt nutrient solution that contains 0.02 ppm Cu (Phytotechnology Laboratories cat. #H353), and stratified in the dark for five days at 4°C. Seeds were then placed in constant 20°C for 14 hour days for 10 days. Seedlings were then transplanted to hydroponic boxes with perforated drinking straws affixed to the cut microcentrifuge tubes (Selby et al. 2014). Plants were transferred to either control conditions (0.5x Hoagland’s solution) or treatment conditions (0.5x Hoagland’s solution with 0.125 ppm CuSO_4_). Solutions were changed every third day. After 18 days, ∼1 inch of the root tips were harvested using forceps then flash frozen in liquid N_2_ and stored at -80°C. RNA was extracted using the Zymo Research Direct-zol RNA kit (cat. #R2070). RNA from one individual per genotype, per treatment, and per replicate was submitted to the Duke Sequencing Facility for library preparation and sequencing. Libraries were made using Illumina TruSeq Stranded-RNA kit, with PolyA enriched mRNA. Samples were pooled and the pool was divided evenly between two lanes of the Illumina HiSeq 2000/2500 platform, generating 50 bp single-end reads.

#### RNA-seq analysis

Libraries were analyzed with FastQC (http://www.bioinformatics.babraham.ac.uk/projects/fastqc/) to assess the quality of the sequences. Reads were mapped to the reference genome *Mimulus guttatus* TOL v5.0 using 2-pass STAR version 2.7.1a (*58*) and raw counts were obtained using --quantMode GeneCounts.

Differential gene expression analysis was done using limma in R Bioconductor, with empirical weights estimated for each observation using the voomWithQualityWeights function (*59*). Quantile normalization was used to account for different RNA inputs and library sizes. The linear model for each gene was specified as: log(counts per million) of a particular gene = genotype + treatment + genotype:treatment. Specific contrasts were constructed to compare each genotype to genotype and each genotype × treatment interaction. Differentially expressed genes were selected based on a false discovery rate < 0.05. DGE pipeline from https://github.com/rodriguezmDNA/rnaseqScripts.

#### Allele specific gene expression analysis

Aligned reads from RNA-seq analysis were used to generate variant call file using the GATK RNAseq short variant discovery (SNPs + Indels) best practices workflow (https://github.com/gatk-workflows/gatk3-4-rnaseq-germline-snps-indels) (*60*). VCF was filtered for gene coding regions for genes within the sweep regions. Allele specific calls were generated following the pipeline developed in (*61*). (https://github.com/hh622/Maize_Highland_Adaptation_allele_specific_expression/tree/main/AS <u>EPipeline</u>). SNP locations were assigned to gene identities. SNPs across the gene were summed (*62*) and an ASE score was calculated for every gene (*63*). Genes with 0 read counts for one of the parental alleles were removed from this analysis. ASE bias was analyzed using a t-test looking for divergence from the expected 0.5 allele balance across the three biological replicates.

#### Genotypes and populations used in this study

##### Inbred lines derived from wild samples

**CCC**. Population derived from Copperopolis Central Court, Copperopolis, CA, a contaminated mine site. Homozygous for *Tol1^CCC^* copper tolerant allele.

**MED84**. Inbred line derived from an uncontaminated inland annual population from Moccasin, CA. Homozygous for a non-tolerant allele *Tol1^MED^* from an inland population.

**STB.** Clonally propagated plant from Stinson Beach, CA, non-tolerant. Used as host genotype for introgression and for mapping populations.

**IM62**. Inbred line derived from Iron Mountain, Oregon, USA (44.402217°N, 122.153317°W). Several versions of the IM62 genome have been released including a preliminary subchromosomal draft sequence IM62 v1.0 (*25*) used for initial genetic mapping of the Tol1 locus and the chromosome-scale reference assembly IM62 v3.1 (*26*) used for subsequent analyses.

**OBR, SCH**. Paired off-mine populations (OBR/CCC) (SCH/MCN)

**MCN**. Population derived from the McNulty Mine in Copperopolis CA, an contaminated mine site.

**MAC.** Population derived from Mackville Road in Dogtown, San Joaquin County CA.

##### Recombinant derived genotypes and lines

**HET_Tol**. Near isogenic line derived by repeated backcrossing of the tolerant CCC line into the intolerant STB line, with genotype *Tol1^CCC^/tol1^STB^* at the Tol1 locus based on tightly linked markers, and homozygous STB elsewhere. This is identical to the genotype 25_E11 discussed in (*23*).

**TOL.** A selfed progeny of NIL 25_E11 that is homozygous for the *Tol1^CCC^* allele based on existing markers linked to the Tol1 locus. The genome of this individual was sequenced and assembled to chromosome-scale, initially as *Mimulus guttatus TOL v3.1* and subsequently improved to *Mimulus guttatus TOL v5*. (The TOL population comprises *Tol1^CCC/CCC^* siblings of TOL.)

**NON_TOL.** A selfed progeny of NIL 25_E11 that is homozygous for the *tol1^STB^* allele based on existing markers linked to the Tol1 locus. The genome of this individual was sequenced and assembled to chromosome-scale *Mimulus guttatus NONTOL v4.0*. (The NONTOL population comprises *Tol1^STB/CSTB^* siblings of TOL.)

##### Mapping populations

Fine mapping of Tol1: one heterozygous BC_7_*^STB^ Tol1^CCC^/tol1^STB^*plant was selected and crossed to a homozygous BC_7_*^STB^ tol1^STB/STB^*sibling to generate a mapping population of 4,340 progeny.

##### Fieldwork populations

Backcross lines were derived by introgressing CCC, MED, and STB Tol1 alleles into STB, yielding *BC_4_^CCC^*, *BC_1_^MED^*, and BC_7_*^STB^*.

---

**Supplemental Figure 1A.**
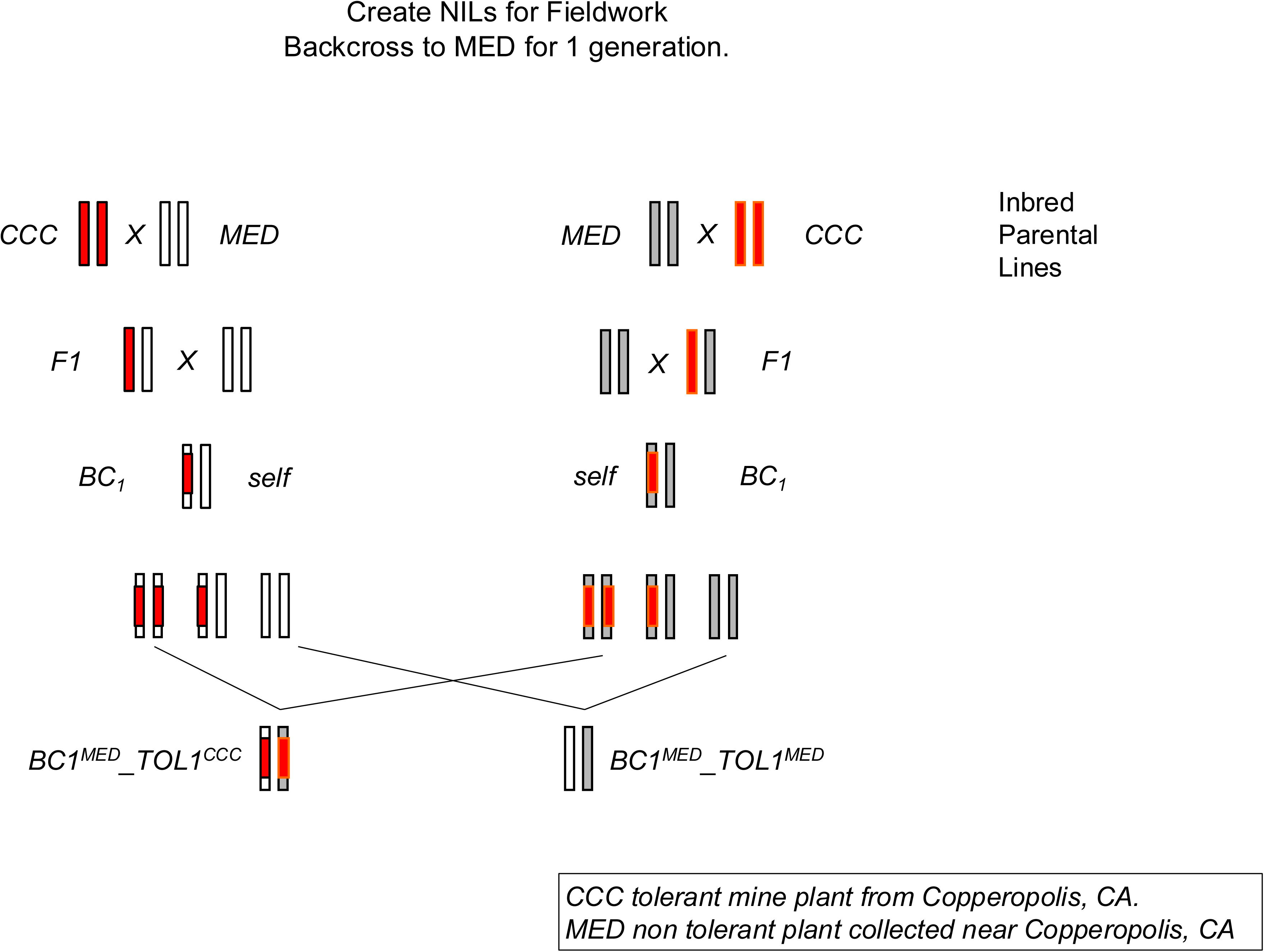

**Supplemental Figure 1B.**
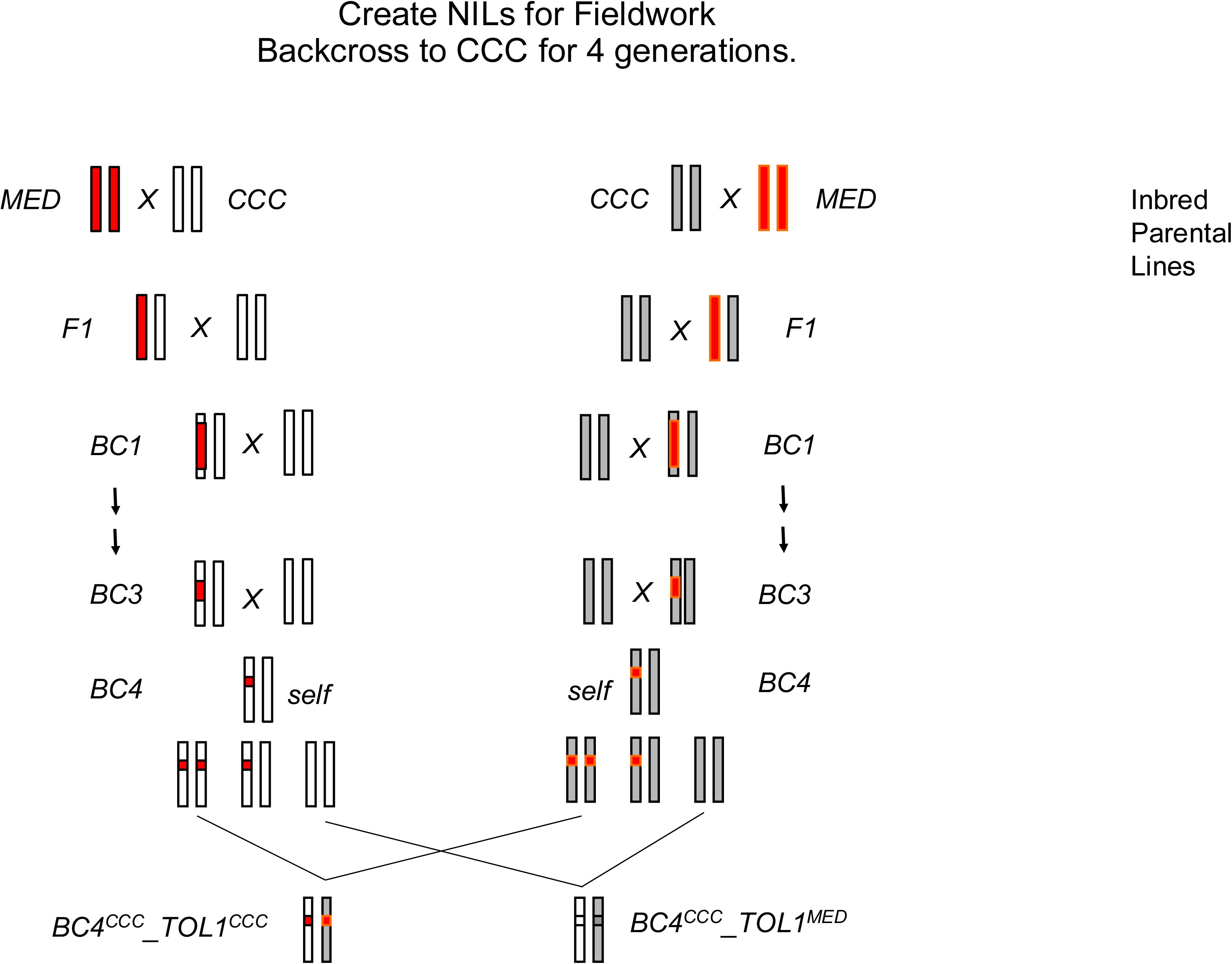

**Supplemental Figure 2.**
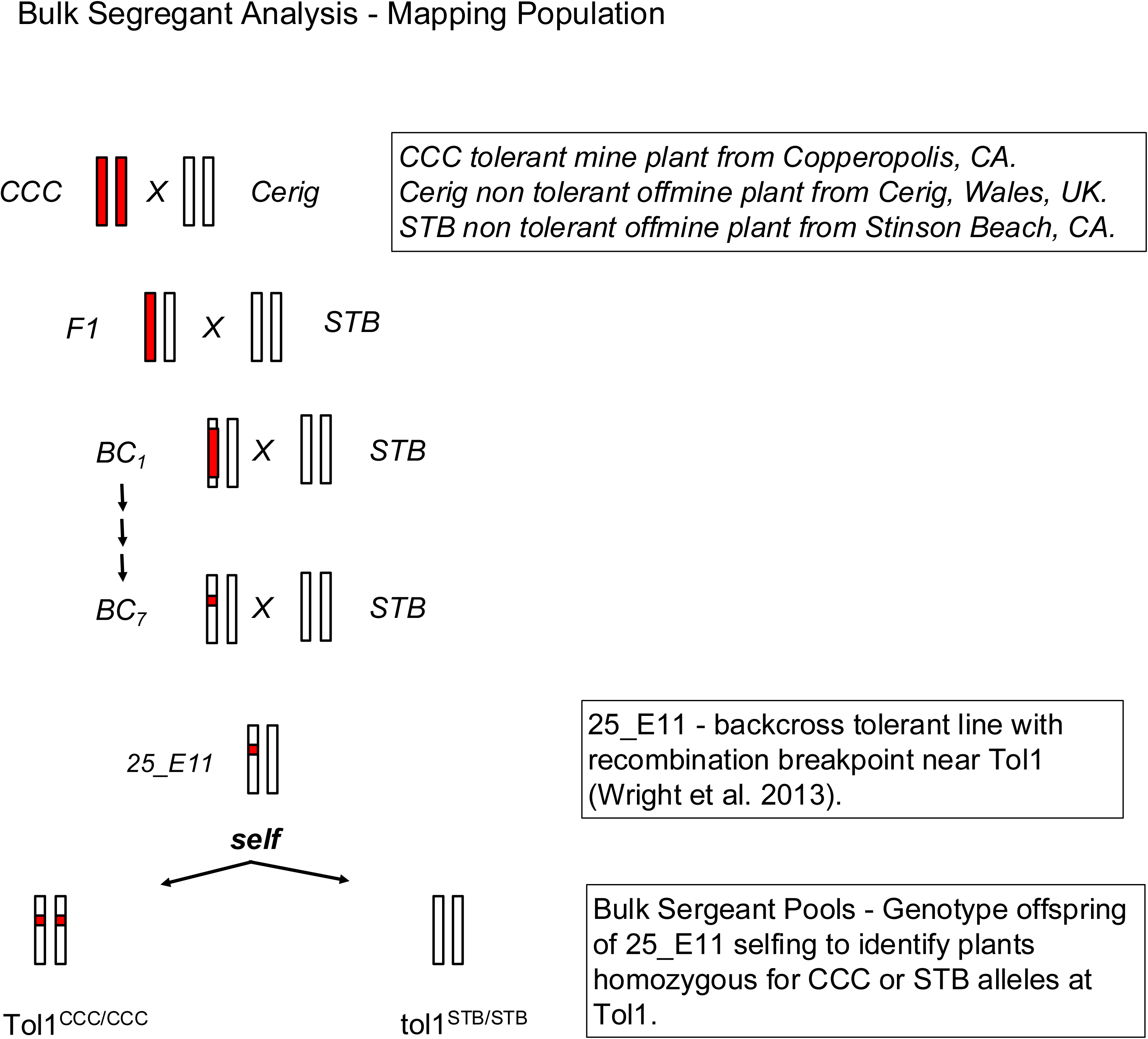

**Supplemental Figure 3.**
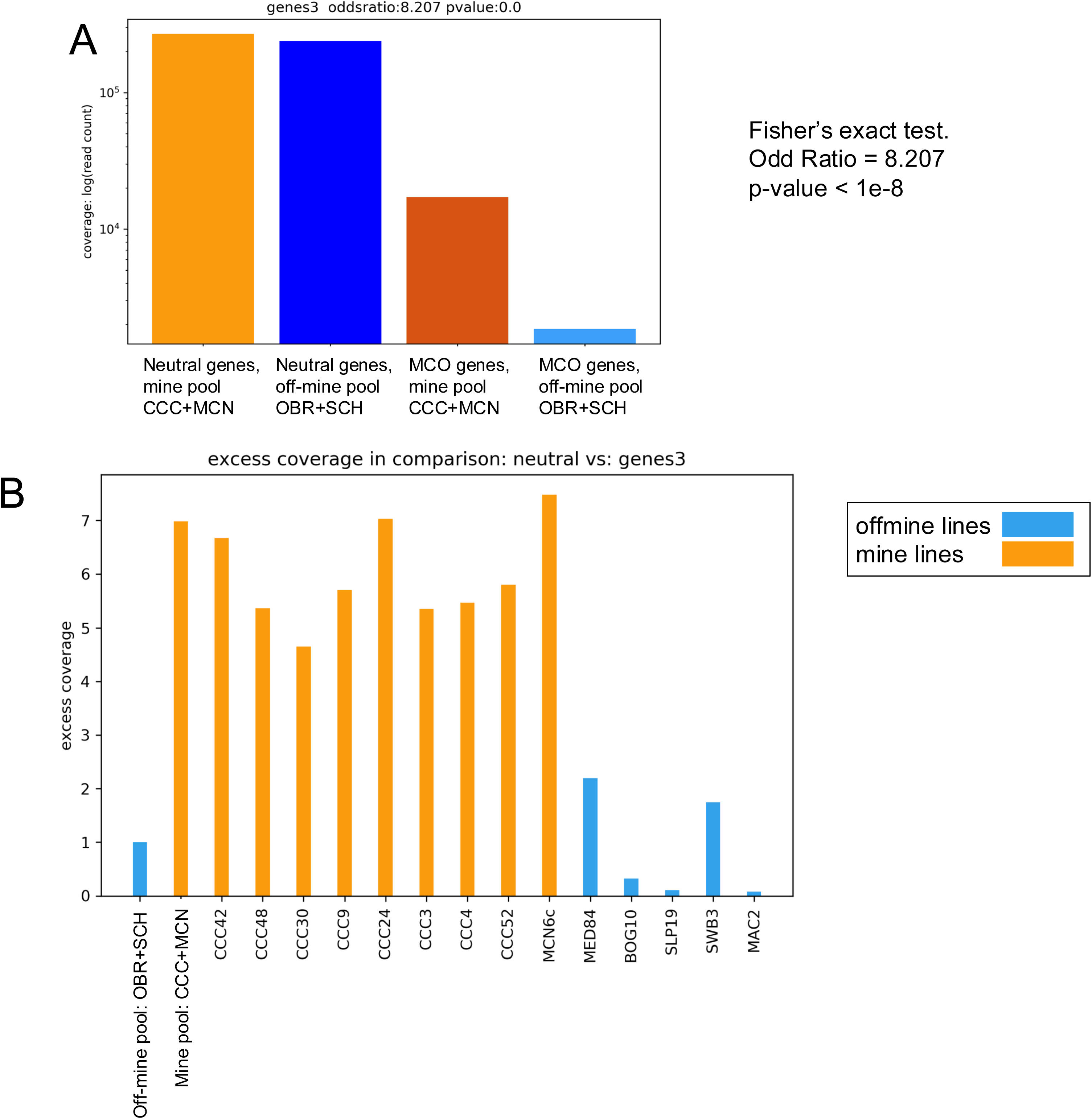

**Supplemental Figure 4.**
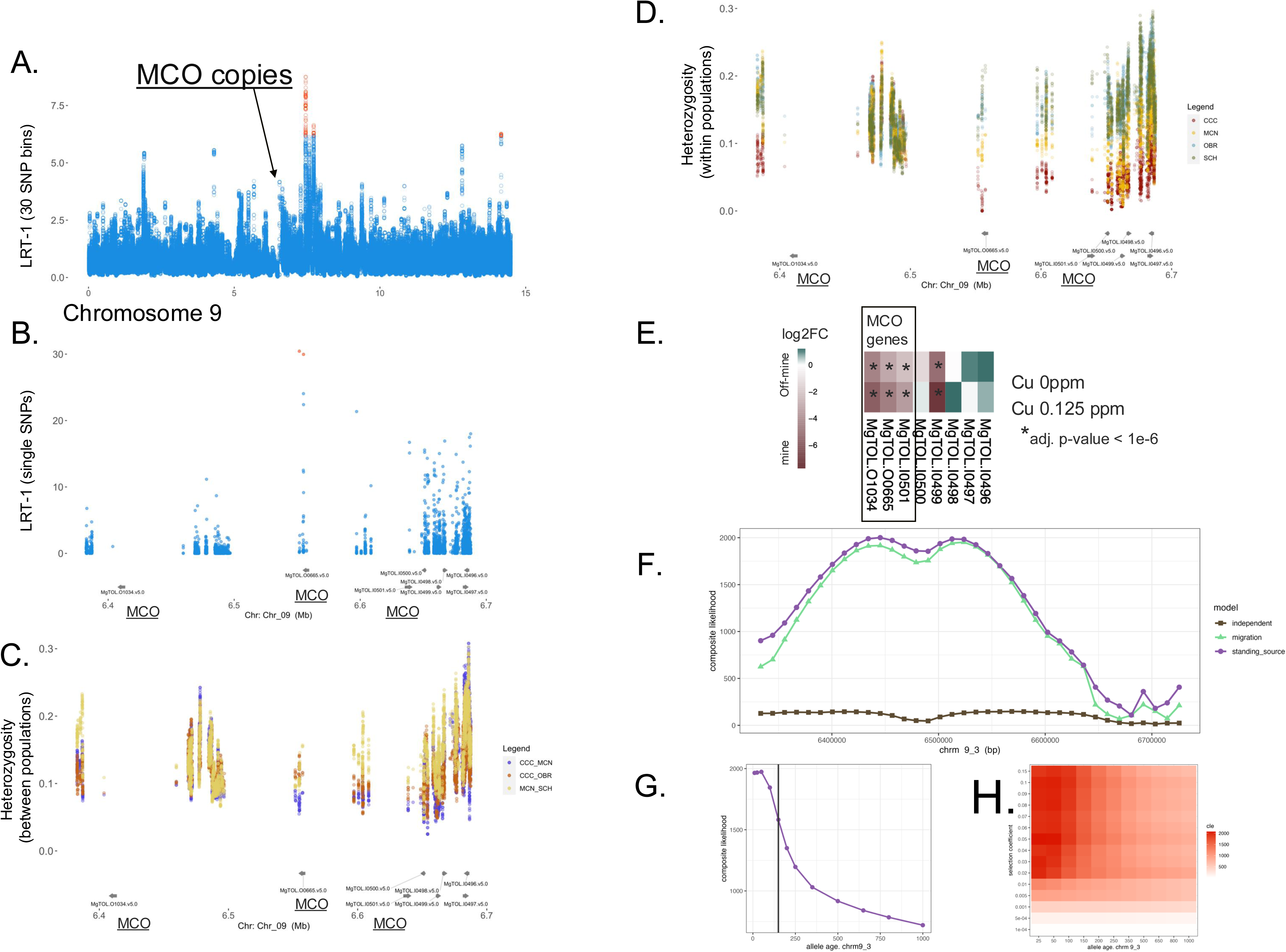

